## Supplemental table description for "Dysregulation in Spatiotemporal Expression of *HMX1* Coordinated by Multifaceted Enhancers Drives an Auricular Disorder" (from another version of this preprint: version 1, 2024-04-02)

### Supplementary Table information

**Supplementary Table 1: PC-HiC interaction information between *HMX1* promoter and other genomic regions.**

Each row in the table contains the genomic locations (hg19) of two regions of the *HMX1* promoter and it's interacting locus, and the interaction score also shows.

**Supplementary Table 2: Primer lists used in this paper.**

The table contains four sheets: RT-PCR primer, LUC primer, LacZ primer and Genotyping primer. RT-PCR primer sheet contains primers for gene expression analysis; LUC primer sheet contains primers of constructing luciferase-related vectors for cell transfection; LacZ primer sheet contains primers for constructing enSERT assay-related vectors; Genotyping primer sheet contains primers for transgenic cell line and mouse models genotyping and validation.

**Supplementary Table 3: Transcription factor binding sites (TFBSs) prediction using FIMO on the hEC1 sequence.**

The table contains motif cluster, motif ID, position, TF symbol, TF class, TF family, score, and *p*-value and q-value for each motif (HOCOMOCO V12 database[1]) bound by TFs that are co-expressed with *HMX1* in craniofacial mesenchyme predicted by FIMO software[2] on the hEC1 sequence. Five clusters of motifs were divided based on the distribution density of motifs, termed as D1 (1~162 bp), D2 (152~292 bp), D3 (282~463 bp), D4 (453~630 bp) and D5 (620~801 bp). The novel coordinator pattern consisting of HD, HMG-box, Coordinator motif in D1 and D3-4 region are highlighted.

**Supplementary Table 4: Results of identified protein binding to hEC1 using LC-MS.**

The table contains protein information identified in the DNA-pull down assay and subsequent LC-MS analysis. And the table is sorted by the score for each protein, and Meis1, Pbx1/3, Dlx6 and Hmga2 TFs were highlighted.

**Supplementary Table 5: EChO analysis results of TCF7L2 CUT&RUN on hCNCCs.**

The table contains information on each position and its corresponding fragment size on the hEC1.

**Supplementary Table 6: Probe sequence information used in EMSA and *in-situ* RNA hybridization.**

DNA sequence information used in EMSA and WISH are provided.

**Supplementary Table 7: Differential gene expression analysis.**

The table provides the information of differential expression genes (DEGs) between $mEC1^{dup/dup}$ and Wild-type (WT) pinna structure at E14.5 mouse embryo. Five columns: Ensembl gene ID, gene symbol, log2 fold change (logFC), mean expression level (logCPM), false discovery rate (FDR), and down- or up-regulated categories (group) in $mEC1^{dup/dup}$ compared with WT. See Methods for details on statistical analysis.

**Supplementary Table 8: GO enrichment analysis.**

The table contains GO enrichment results of each GO ID, description, GeneRatio, Background ratio (BgRatio), p value, p.adjust (adjusted by Benjamini-Hochberg, BH), q value, geneID and gene count (Count).
